# Immune Antibodies Recognizing the Stem Region of SARS-CoV-2 Spike Protein: Molecular Modelling and *In Vitro* Study of Synthetic Peptides Presentation to the Antibodies

**DOI:** 10.1101/2025.03.20.644166

**Authors:** Elena T. Aliper, Ivan M. Ryzhov, Polina S. Obukhova, Alexander B. Tuzikov, Oxana E. Galanina, Marina M. Ziganshina, Gennady T. Sukhikh, Nikolay A. Krylov, Stephen M. Henry, Roman G. Efremov, Nicolai V. Bovin

## Abstract

Antibodies to peptide 1147 (amino acids 1147-61) of the SARS-CoV-2 protein S are highly diagnostic. Peptide 1147, although located in a region that is partly spatially hidden in the intact protein, is not subject to mutations, suggesting therapeutic potential. The aim of this study was to elucidate the architecture of this region and the way in which it is presented to antibodies. As a model system, this peptide carrying a single lipophilic tail and the same peptide carrying a lipophilic tail at both ends (pseudocyclic) were incorporated into lipid membrane. Isolated anti-1147 antibodies interacted with it regardless of how the peptide was presented, be that freely exposed via the N-terminus, organized as a pseudocycle, or adsorbed on the surface. MDS showed that peptide 1147 is capable of closely approaching the membrane. Analysis of the surface properties of peptide 1147 in membrane-bound states and in available conformations in the full-sized S protein reveals interface for interaction with antibodies. Interestingly, the latter bears similarities to one published peptide-antibody complex. However, these antibodies, in spite of their high diagnostic significance, show no virus-neutralizing activity, indicating that peptide 1147 has no therapeutic value as a synthetic vaccine.

## 1. Introduction

Pinpointing the exact epitope specificity of anti-SARS-CoV-2 antibodies conferring therapeutic value on convalescent plasma is a major challenge. Two aspects can be distinguished here: one is associated with the phenomenon of antibody-dependent enhancement (ADE) [1], while the other is the necessity to identify the most effective plasma antibodies. On this basis, one would be able to develop monoclonal antibodies with similar specificity. In contrast, selecting epitopes suitable for diagnostic antibody screening, although it focuses on finding conservative epitopes, does not require for these epitopes to possess therapeutic value. In fact, it is well known that many diagnostic (and highly immunogenic) epitopes only become exposed after viral degradation and may be poorly exposed in the intact protein/virus. When searching for the S protein’s diagnostic epitopes suitable for plasma screening of individuals who have had COVID-19, five peptides of the spike protein were initially identified and then used to build conjugates, lipophilic function-spacer-lipid constructs (FSLs). These peptide-based FSL constructs were inserted into the human red blood cell membrane, resulting in a novel diagnostic assay [2-4]. Using these peptidated cells as a diagnostic system, a cohort of convalescent COVID-19 patients were examined for high titers of therapeutically significant antibodies [2]. Only one of the peptides predicted initially, SFKEELDKYFKNHTS, named peptide 1147 after the position of its first residue in the amino acid sequence of the S protein (Fig. 1) showed both high specificity and sensitivity in detecting antibodies (IgG), as well as correlation of antibody level to disease severity.

**Figure 1.**
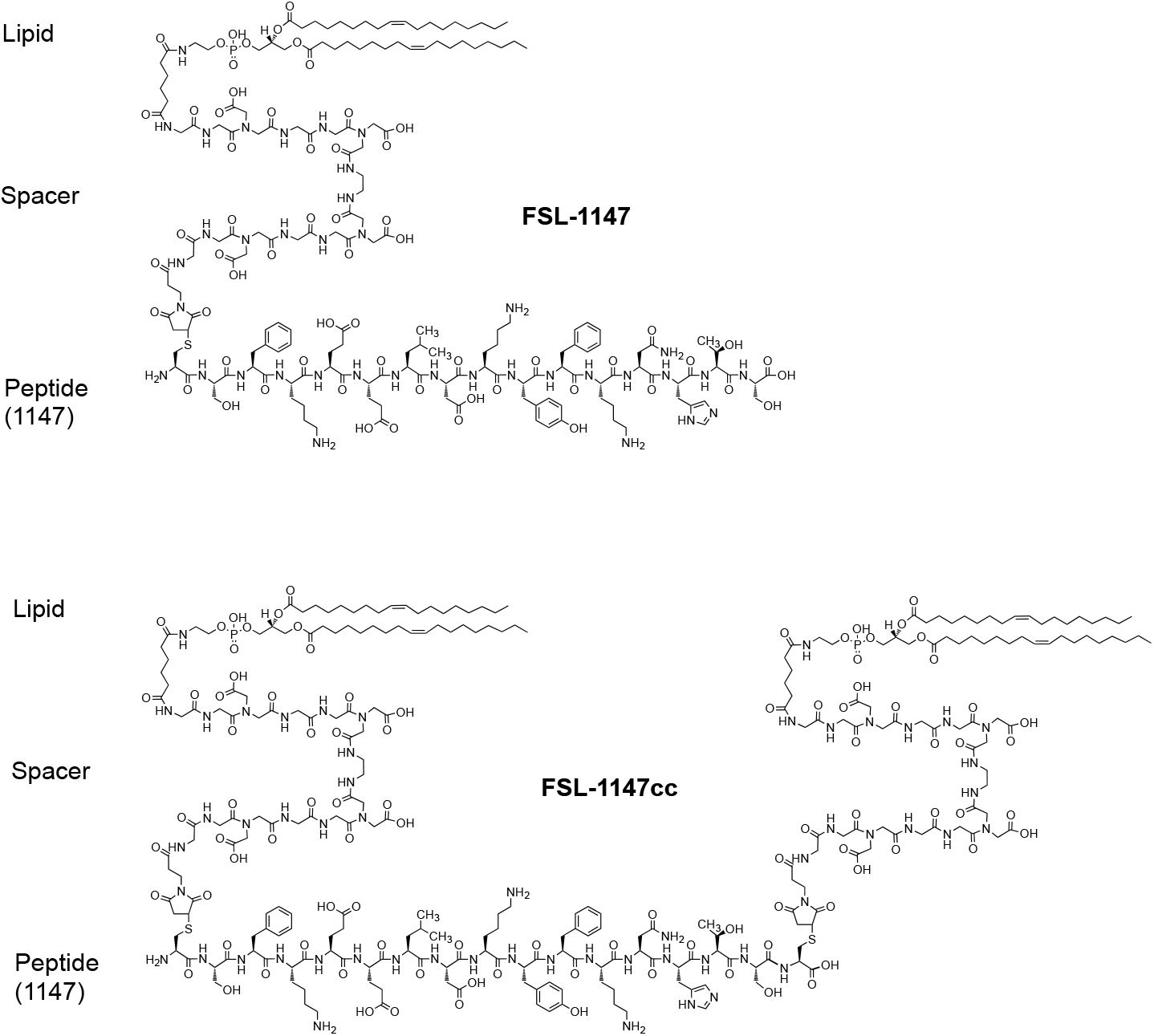
Structure of the mono-lipidated linear FSL-1147 and the dual-lipidated pseudocyclic FSL-1147cc constructs, the latter with two DOPE residues attached via GMG spacer to both the C- and N-termini. Peptide 1147 with the amino acid sequence SFKEELDKYFKNHTS in both constructs is conjugated to the CMG spacers via an additional terminal cysteine residue.

Intriguingly, in the pre-fusion state peptide 1147 is not sufficiently far from the lipid membrane in which the S protein is anchored (Fig. 2); this implies potential difficulties for recognition by antibodies.

**Figure 2.**
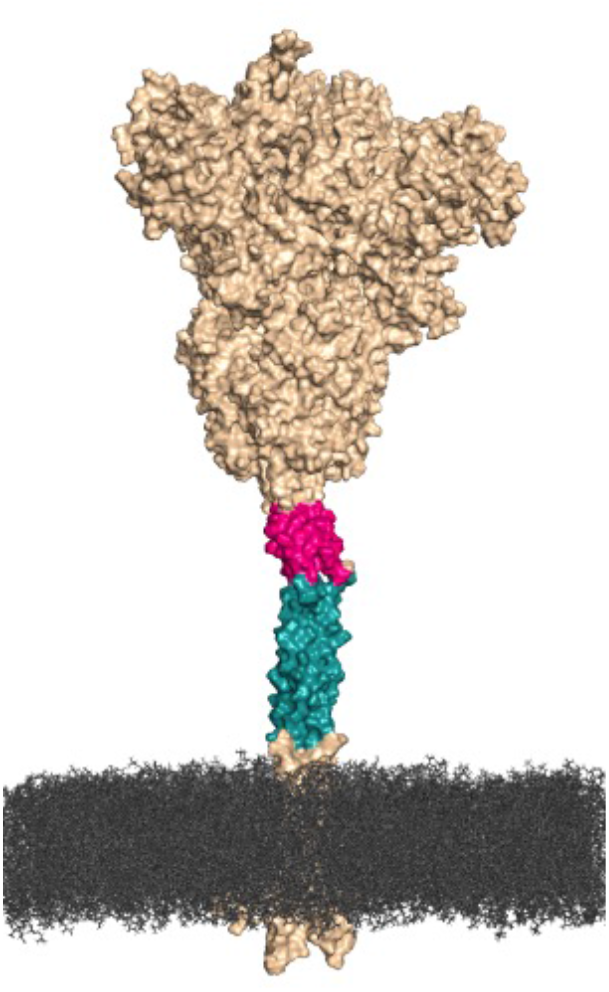
Peptide 1147 (amino acid residues 1147-61), is located in the pre-fusion spike (colored magenta) immediately upstream of the functionally crucial HR2 region (colored green) of the S protein.

We therefore put forward several hypotheses, taking this into account along with certain experimental data

1. Anti-1147 antibodies can only be generated after the degradation of the virus and the S protein, whereupon the highly immunogenic 1147 peptide becomes exposed to the immune system, and antibodies thereto start developing. Anti-1147 antibodies are therefore a consequence of viral destruction/degradation and are probably not actively involved in virus neutralization, although they may be an S protein scavenger, acting after the destruction of the native virus.
2. Anti-1147 antibodies are able to bind the intact virus and block the infection. The conservatism of the epitope and its proximity to the membrane suggest their potential for broadly neutralizing properties.
3. Epitopes on peptide 1147 evolved in the SARS-CoV-2 virus as a feature counteracting the host’s immune system via the ADE mechanism [1], discussing which in detail would go beyond the scope of this article

To examine these hypotheses, it is necessary to know the architecture, structural, amphiphilic, and dynamic properties of that part of the S protein stem, especially the peptide 1147 region, as well as what happens during its immunological presentation as an antigen. To put these speculations to test, molecular modeling was used, and convalescent’s antibodies specific to peptide 1147 were isolated; their interaction with peptide 1147 under conditions simulating significantly different modes of its presentation, as well as virus neutralizing potency, was studied.

## 2. Results and Discussion

### Synthesis of FSL-1147cc and other FSL constructs

Most experimental work was carried out on the 16-mer peptide 1147, [C]SFKEELDKYFKNHTS, which includes an additional cysteine residue (for conjugating, designated [C]), although some of the experiments were performed on the 16-mer peptide 1144, [C]ELDSFKEELDKYFKN. The FSL based on peptide 1144 is similar to the peptide 1147-based one, however, it is shifted 3 residues upstream along the amino acid sequence of the S protein. Sensitivity and specificity comparison of these two different peptides as FSLs showed a small improvement in performance for peptide 1144, and no significant difference when interacting with patient-*vs*-donor antibodies (data not shown). Synthesis of monolipidated FSLs such as those based on peptide 1147 (Fig, 1) and peptide 1144 have been described previously [5]. In contrast, the FSL-1147cc construct which has two lipophilic anchors was synthesized via conjugation of the peptide [C]SFKEELDKYFKNHTS[C] (with cysteine residues at both ends) with two moles of spacer-DOPE also comprising a maleimide group. The spacer designated as CMG(2) is assembled from a triglycine block, where one of the glycine residues has a nitrogen atom modified with a -CH_2_-COOH group [6]. This design of the FSL-1147cc construct, due to its two lipid tails, would be expected to become embedded in a lipid layer at both ends, while the negatively charged and rather rigid but centrally hinged CMG(2) spacers would ensure that the peptide fragment would not directly contact the artificial lipid membrane, or the plasma membrane in the case of live virus entry into cells. Synthesis protocols have been described previously [7].

### Affinity isolation of anti-1147 antibodies

Isolation of immune antibodies is described in detail elsewhere [5]. In short, peptide 1147 was attached to SepharoseFF resin, and a pool of sera from patients recently recovered from Covid-19 and previously identified as having a high titer of the corresponding antibodies was passed through it. Antibodies were gently eluted from the column and then depleted on peptide-free SepharoseFF to eliminate the antibodies to the matrix, which are always present (up to 30%) when this method is applied [8] and could influence subsequent experiments.

### Binding of affinity-isolated antibodies to differently presented peptide 1147

The peptide 1147 SepharoseFF affinity-isolated antibodies were tested against linear peptide 1147, pseudocyclic peptide 1147cc (see structures in Fig. 1) and peptide 802 (as a negative control). The FSL constructs were incorporated into a lipid layer pre-formed of PC and cholesterol in the microwells of EIA (enzyme immunoassay) polystyrene plates using two different approaches (briefly described elsewhere [5]). The first approach only includes one step: applying a solution of PC, cholesterol and FSL (in which the proportion of FSL varies) in ethanol onto the plastic. The second, two-step approach consists in the preparation of a PC-cholesterol layer followed by the incorporation of the FSL (in varying concentrations) from an aqueous solution to the pre-existing lipid layer. In addition, all three constructs (as well as FSL with PEG instead of a peptide as an additional negative control) were analyzed using EIA.

The results (Fig. 3) of anti-1147’s interaction with the FSL variants indicate an almost identical mode of binding with linear peptide 1147 and the pseudocyclic variant 1147cc regardless of whether the FSL was incorporated into a pre-formed lipid layer in an orderly manner (two-step approach) or whether an FSL/lipid layer was prepared from a mixture of lipids including the FSL (the one-step method). It is of note that our results are not in agreement with an independent report [9] stating that the author’s cyclic epitope 1146 interacted better with antibodies from Covid-19 patients compared to the linear variant; however, in this study the peptides were not presented to the antibodies in the lipidated form described in this paper.

**Figure 3.**
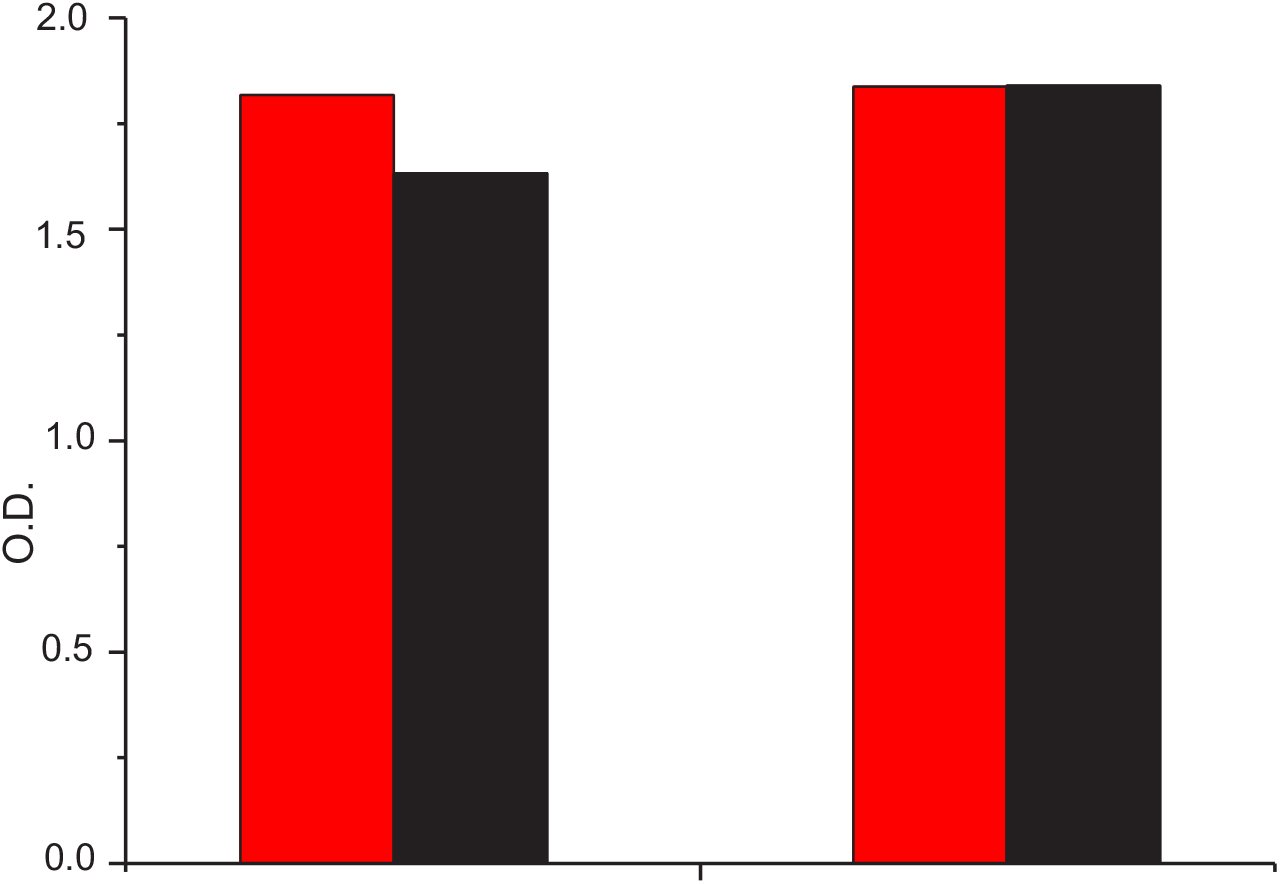
Binding of affinity isolated anti-1147 antibodies to differently presented 1147 peptides, EIA data. Left pair: FSL-1147cc (red bar) and FSL-1147 (black bar) and were inserted into a PC/ cholesterol layer using the one-step method at an FSL/PC molar ratio of 0.087. Right pair: FSL-1147cc (red bar) and FSL-1147 (black bar) were inserted into the lipid layer using the two-step procedure at 39 ng of FSL added per well. In both cases, the maximum values of the OD of the corresponding concentration plots are presented.

### Influence of the cell membrane environment on the binding of antibodies to peptide 1147

The inaccessibility of the peptide region 1147-1161 in the intact virus to antibodies can be explained by both architectural (or conformational) restrictions in the S protein trimer (data on the simulation of this fragment’s interaction with the membrane are presented below), as well as by the natural environment of the protein when located in the live virus membrane. We evaluated the latter experimentally via the insertion of FSL-1147 into an erythrocyte membrane (although not the same as a virus membrane, it would at least simulate restrictions imposed by the lipid membrane, which might impact how antibodies approach the peptide). We studied its interaction with antibodies using flow cytometry (by detecting anti-1147 with fluorophore-labeled antibodies to IgG). This experiment did not use affinity-purified anti-1147 (discussed above); instead, pooled serum from patients with Covid-19 was used. The results (Supplementary Data, Fig. S6) show that antibodies to peptide 1147 in the serum bind to this antigen (in the form of FSL) inserted in the cell membrane.

### Virus-neutralizing potency

The potential antiviral activity of affinity-isolated antibodies to peptide 1147 was examined by testing their ability to prevent live virus from infecting a Vero C 1008 cell culture [1], and comparing results to those obtained for convalescent serum (the source used for affinity isolation of anti-1147). The serum which would be expected to contain several different antibodies to the S protein (and other viral epitopes) 100% blocked the activity of the virus. In contrast, the efficiency of affinity-isolated monospecific antibodies in blocking the virus was much lower, only reaching 10% at high concentrations (>1 μg/ml, Fig. 4).

**Figure 4.**
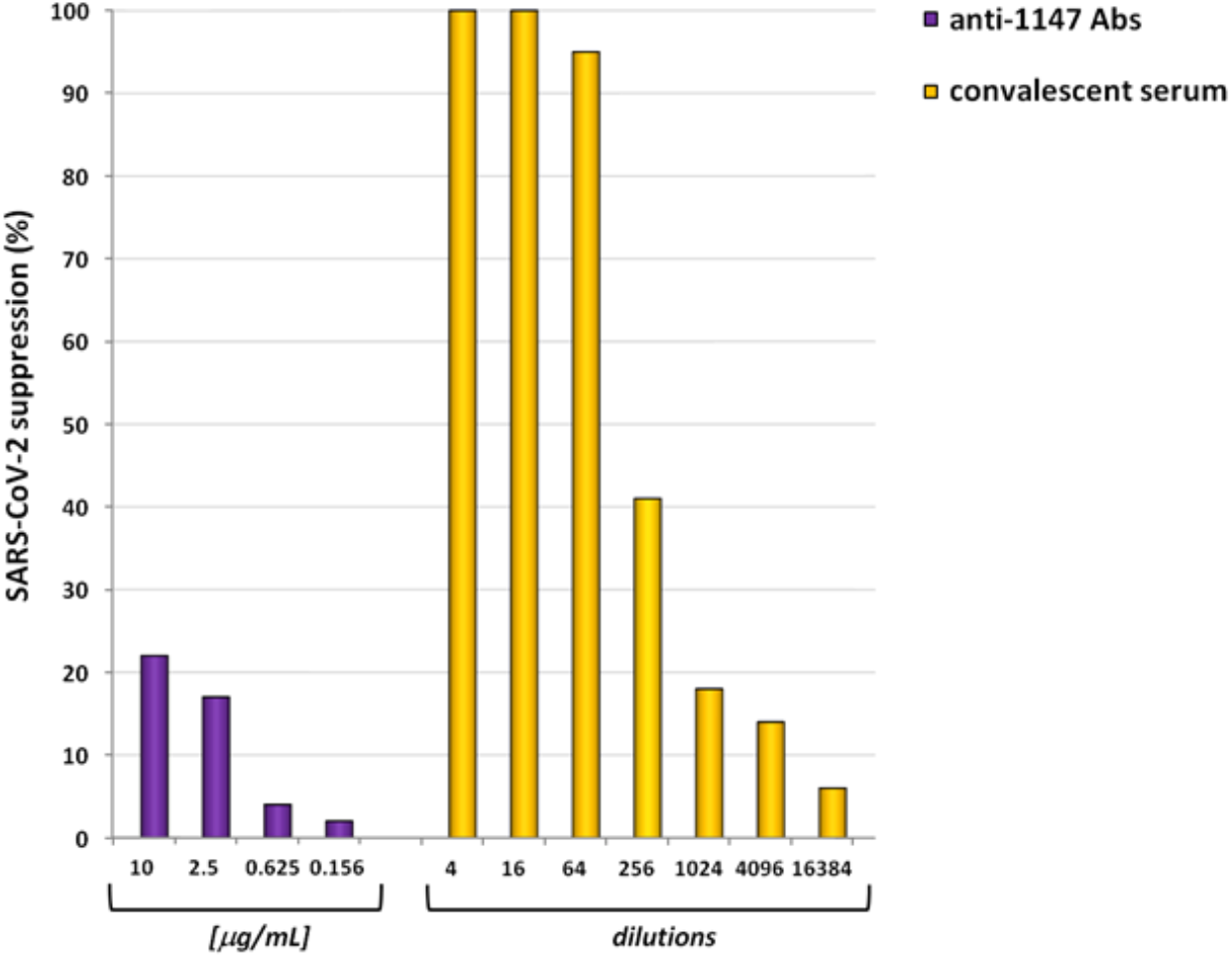
Convalescent serum neutralization of SARS-CoV-2 virus in the Vero C 1008 cell culture experimental system.

### Design of the molecular modeling study

A modelling-based approach was employed that included the following steps: 1) Detailed mapping of the object’s surface regions for two states known for the spike homotrimer, being pre-fusion (PDB ID 6XR8) and post-fusion (PDB ID 8FDW). 2) A model of the FSL-1147 (mono lipidated, see structure in Fig. 1) for which MD simulations were run in an artificial lipid bilayer to identify any binding modes. Peptide 1147 behaved differently across several trajectories calculated. Although there were several trajectories in which the peptide bound transiently or poorly, of interest to us were only those states in which binding took place in a membrane, since the primary goal was to establish which residues might be accessible to the antibody. Two such states were found, one in which the peptide was immersed deeply in the lipid bilayer and the other in which it was superficially embedded; both of these states were further studied.

Surface physicochemical properties of peptide 1147 were calculated in each of four states (pre-fusion, post-fusion, tightly bound FSL, loosely bound FSL), along with aqueous solvent-exposed and buried residues. 3) A consensus characteristic pattern of residues accessible to the aqueous solvent in all four states was identified. 4) To test whether the peptide 1147 with delineated pattern of solvent-exposed residues can bind to Ab, we performed a database search of known spatial structures for complexes containing antibodies with peptides/proteins carrying the motif identified at step 3. One high-scoring match was found, and it was shown that its physicochemical properties and interaction with a cognate antibody agree well with the available data on peptide 1147. These results were used to elaborate a model of the potential Ab-binding site and propose its further modifications, which can affect epitope 1147/Ab interaction.

### Structural analysis and molecular dynamics simulation

In the pre-fusion spike peptide 1147 is α-helical and essentially amphiphilic, with a distinct hydrophobic patch on one of the surfaces (made up by residues F1148, L1152, Y1155 and F1156, and is either identical or conserved across beta-coronaviruses, as analyzed using data from UniProt [11]). Peptide 1147 is packed like a canonical water-soluble coiled-coil with the hydrophobic patch hidden away inside the trimer lumen. In the post-fusion state, the hydrophobic residues are inaccessible to polar solvents, although the α-helix becomes partially disordered at the C-terminus of the peptide (Supplementary data, Fig. S1) and forms an interface with the trimeric central helix domain (CH) [12]. Fewer residues remain solvent-accessible compared to the pre-fusion state (see Fig. 5 and Fig. 7).

**Figure 5.**
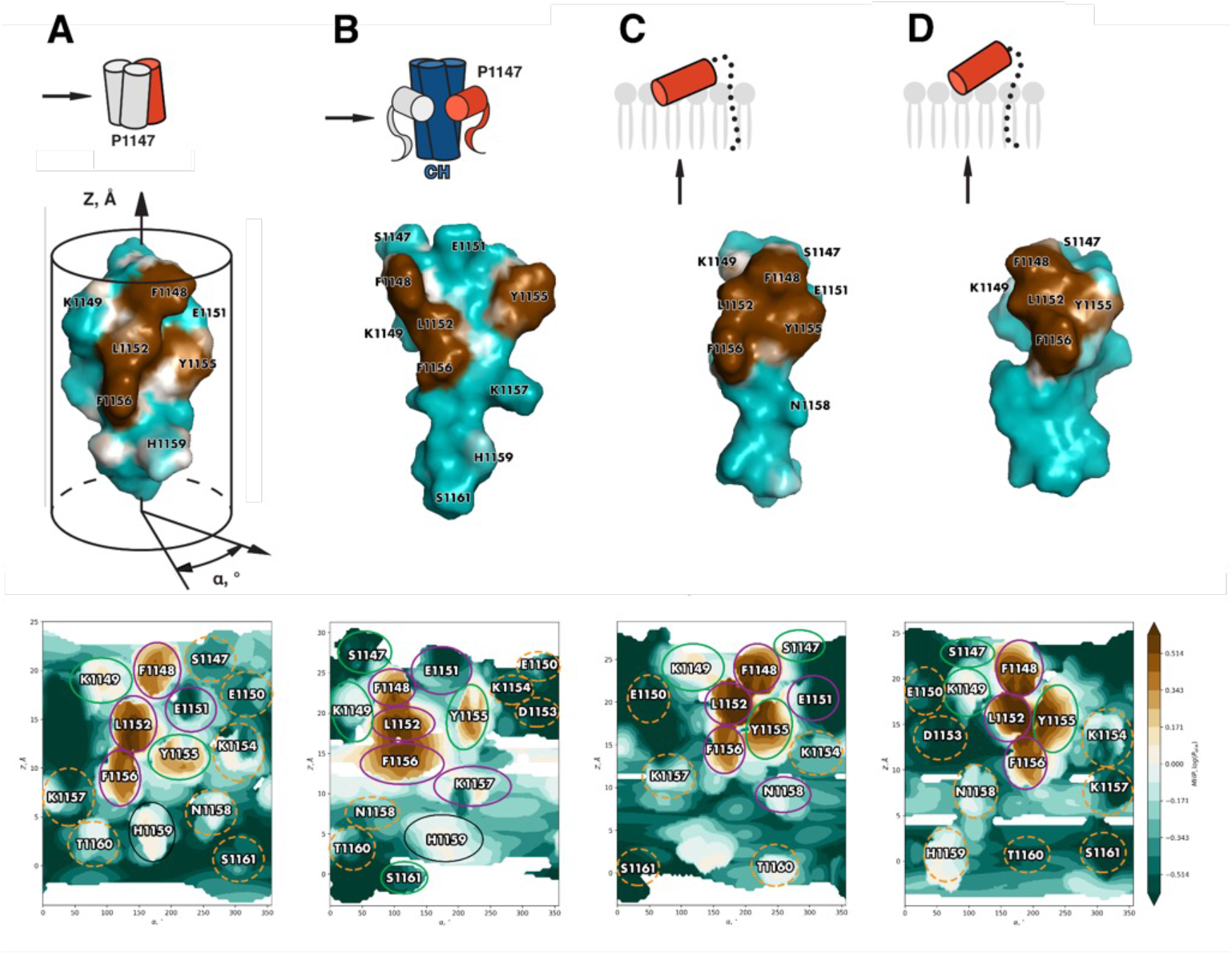
Physicochemical properties of peptide 1147 in different states. (A) as part of the pre-fusion spike, (B) as part of the post-fusion spike bound to the central helix (CH) domain, (C) in tightly, and (D) in loosely bound states in a model membrane as determined by MD of the FSL. The top row schematically shows the localization of peptide 1147 in various states (the spacer and lipid of the FSL are represented as dotted lines), while the arrows indicate the angle from which the object, colored red (shown in the middle row), is viewed. The middle row shows 3D models of the same states in surface representation colored in accordance with MHP values (hydrophobic and hydrophilic zones are colored brown and teal, respectively). The 3D model for the pre-fusion state is enclosed in a schematic cylinder completed with axes used to plot the 2D maps in the bottom row. The bottom row shows 2D cylindrical projections of surface MHP distribution. Axis values correspond to the rotation angle around the helical axis and the coordinate along the latter (Å), respectively. A MHP scale (in logP octanol-1/water units) is presented on the right. The maps are colored in accordance with the MHP values,^[10]^ from teal (hydrophilic areas) to brown (hydrophobic ones). Of the residues inaccessible to the solvent, identical positions are encircled in purple, conservative and semi-conservative residues are encircled in green, non-conservative residues present on the helix/helix interface are encircled in black. Meanwhile, solvent-accessible residues are enclosed in dashed golden contours.

**Figure 6.**
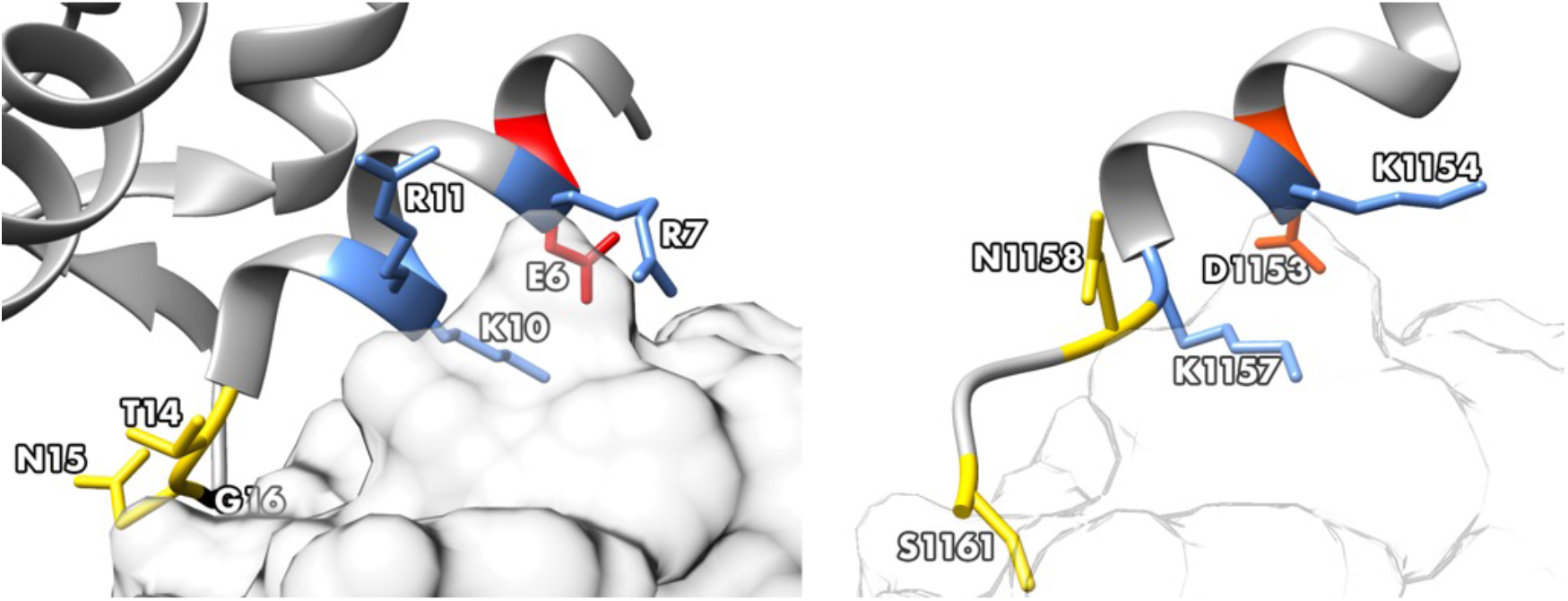
Comparison between the spatial structure of *Phl p 7* noncanonical epitope (left) and peptide 1147 (right). The peptides are shown in cartoon representation, while the antibody binding *Phl p 7* and a hypothetical outline of the same projected against peptide 1147 are shown in surface representation. Residues forming the interface with the antibody in *Phl p 7* and their color-matched counterparts in peptide 1147 are shown in stick representation. Cationic (blue), anionic (red), polar (yellow) and small residue (black) interfaces with the antibody are shown.

**Figure 7.**
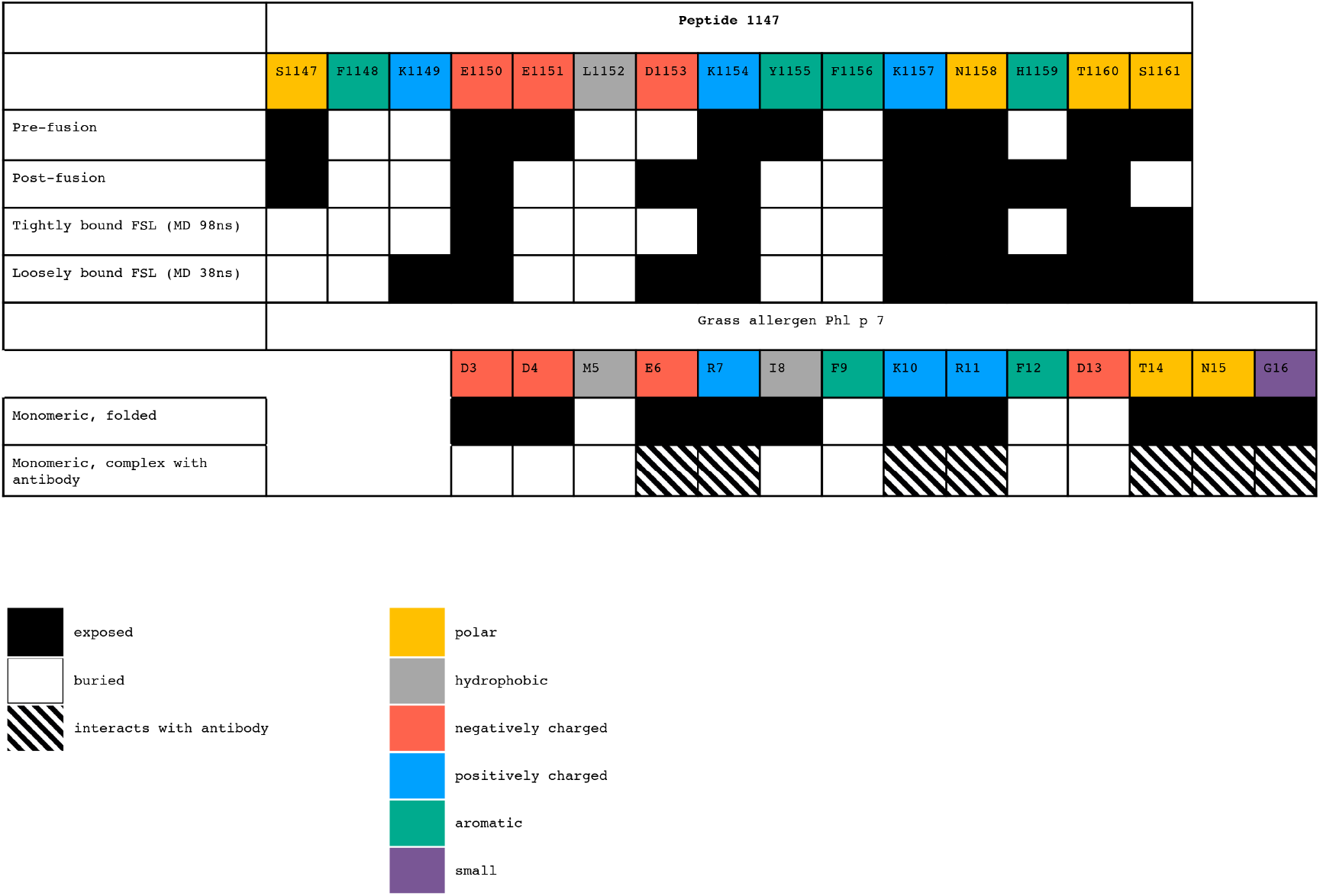
Accessibility to the solvent of amino acid residues in peptide 1147 and grass allergen Phl p 7.

It has been speculated that during refolding and membrane fusion this particular portion of the spike essentially operates like a monomer. This hypothetical monomeric state of anchoring close to the membrane corresponds to a membrane modelling of the FSL (Fig. 5). Accordingly, we modeled a system with an FSL molecule anchored in a POPC/Chol bilayer (2:1), and MD simulations revealed a long-lived membrane-bound state for peptide 1147. The peptide remains mostly α-helical and is only partially disordered at the C-terminus, thus mostly preserving its amphipathic nature. It also appears tilted with respect to the bilayer surface, as is often the case with membrane-active peptides, with the hydrophobic residues F1148 and K1149 buried the most deeply. The mobility of the FSL moieties, the geometry of the membrane binding mode, as well as non-covalent peptide/lipid interactions were analyzed in detail (see Supplementary data). Data suggest that the membrane binding mode is energetically favorable, which explains peptide 1147’s firm attachment to the membrane. However, for such tight membrane binding to happen, a loosely bound state is also required, with a higher solvent accessibility of the residues. In both cases, the interaction pattern in the membrane-bound state is indicative of the way the peptide 1147 is packed when part of the complete spike protein, that is the hydrophobic patch seek to escape the aqueous environment by interacting with lipids, like it does in the pre-fusion and post-fusion states of the spike (see Fig. 5 and Fig. 7).

Based on these data on partially/fully membrane buried and solvent-accessible residues, one can speculate that at least some of peptide 1147 must be as an epitope recognized by human anti-1147 antibody. To test this assumption, we independently searched for similar epitopes across the known spatial structures of antibody complexes with peptides/proteins carrying a sequence motif of residues available to the antibody similar to the one found in peptide 1147 (see next section and Methods). These results are summarized below.

### Pattern matching against known structures and epitope prediction

Searches across the PDB database yielded one particularly well-scoring match (PDB ID 5OTJ), where an antibody complexed with the peptide polcacin Phl p7 [13] containing the sequence 3-DDMERIF-9 (Supplementary data, Fig. S5 C), highly similar in physicochemical properties to peptide 1147. The *Phl p 7* antibody-bound helix (helix 1) is distinctly amphiphilic, with a hydrophobic patch parallel to the helical axis and hydrophilic residues on the remaining molecular surface (Supplementary data, Fig. S5 D). The hydrophobic patch in helix 1 faces the remainder of the molecule, while the charged Glu6, Arg7, Lys10 interact with the antibody. In addition, Glu6 forms a salt bridge with an Arg residue in the antibody (see Fig. S5 E, Supplementary data). A very similar surface pattern is also present in peptide 1147, represented by Asp1153, Lys1154, and Lys1157 (see Fig. S5 F, Supplementary data). The *Phl p 7* Asp3 residue does not interact with the antibody which corresponds to Glu1151 in peptide 1147. This residue might be poorly accessible to the antibody due to its proximity to the loop formed by the fragment between the DOPE and peptide 1147 that remains in water and in close proximity to the peptide and could be similarly unessential to antibody binding. The similarities extend to other motifs: the counterpart of Arg11 in *Phl p 7* would be the hydrophilic N1158 in peptide 1147, which, although uncharged, is no longer part of the helix and is located slightly differently in relation to the remainder of the “ionic islet”. Yet, evidence from Fig. 7 and the heatmap in Fig. S5 A (Supplementary data) shows there are periods across the trajectory when N1158 is spatially available to be bound, along with the two lysines and aspartate.

Downstream of F12 the region corresponding to helix 1in *Phl p 7*, it is no longer helical, but disordered, rather like peptide 1147 in our MD simulations downstream of Lys1157, hence we draw no further parallels from this position onwards. However, certain residues (Asp13, Asn15, Gly16) in this disordered segment do interact with the antibody, so must also contribute to antibody interaction, or at the very least be spatially available to do so. On the basis of solvent-accessible surface data these would be residues His1159, Thr1160 and Ser1161 (Fig. 7 and 3D image in Fig. S5 B, Supplementary data).

The design approach taken to develop SARS-CoV-2 antibody diagnostics [2] was to first identify potential non-glycosylated epitopes on the spike protein using a range of predictive algorithms and online tools. The selected peptides were then converted in a lipid-conjugated form and attached to cells where these peptide-loaded cells were used to study antibodies in Covid-19 patients. Peptide 1147 results demonstrated good correlation with the disease and a lack of interaction of samples from healthy donors, good diagnostic sensitivity and specificity. In the previous diagnostic use publication [2], we did not explore the topology of the antigenic determinants, the topic of this article. Initially a number of peptides found in different regions of the spike protein were selected according to their predicted antigenic potential, and the most diagnostically useful was peptide 1147 which was located in the “lower” stem part of the protein S and in close proximity to the lipid bilayer of the virus membrane. From general considerations, it is expected that the target for virus-blocking antibodies should be readily accessible on the intact virus, and those antibodies formed only as a consequence of viral degradation, although of high diagnostic value will have no or minimal therapeutic potency. However, interaction of antibodies with a poorly accessible normally hidden peptide 1147 region may have importance for neutralizing the virus through a mechanism of action on a conformationally mobile fragment of protein S. This assumption is well-supported by the literature [14, 15] where the ability of antibodies (“broadly neutralizing”) to bind similar normally hidden sites of other viral proteins – trimers that function as a fusion factor. The most investigated in this aspect is influenza virus hemagglutinin [16], where it has been found that the stem-directed antibodies play a blocking role in influenza immunity.

Molecular modeling, including MD simulations in a POPC/Chol membrane, revealed that the assumption that region 1147 is inaccessible for recognition by antibodies is not supported. Instead, MD reveals a membrane-binding mode for peptide 1147 within the pre-fusion state, whereby the hydrophobic residues chiefly made up by the bulky F1148, K1149, L1152, Y1155 and F1156 are buried in the membrane. Interestingly, the recently published structure of the spike in the post-fusion state (PDB ID 8FDW) [12] demonstrates a somewhat similar behavior. In that case, the helical conformation is preserved for residues 1148 to 1152, while residues 1153 to 1155 constitute a turn-like structure, and further downstream the backbone adopts an extended conformation. Residues F1148, L1152 and F1156 which face the core of the post-fusion spike trimer, contact the central helix domain (circa res. 990-996), while Y1155 contacts the three-helix (3H) domain positioned nearby (circa residue 755). These more hydrophilic residues thus face away from the core of the spike. One might therefore speculate that the amino acid sequence of peptide 1147 confers upon it the capacity for interacting with hydrophobic/hydrophilic interfaces in a variety of scenarios, some of which are functionally important to the protein at its various stages in its lifecycle. Searches for known complexes of antibodies with proteins containing the [ED][ED]x[ED][KR]xx motif present in peptide 1147 revealed a match with the *Phl p 7* protein which has an ionic islet formed by one negatively charged and three positively charged residues (corresponding to D1153, K1154, K1157, and the polar N1158 in peptide 1147) on the surface of a helix. These residues could be part of an epitope, albeit a “noncanonical” one [13], recognized by antibody, and suggesting that the membrane-binding mode identified in the MD simulations of peptide 1147 may be compatible with recognition by an antibody. Admittedly, N1158 is on the interface with the membrane (see Fig. S5 A, Supplementary data) and not in the helical conformation (Fig. S4, Supplementary data) in the majority of MD states. However, over the course of MD, we were able to detect transient states in which N1158 was solvent-accessible, and this ionic/polar islet in peptide 1147 was extremely similar to the one in *Phl p 7* in terms of spatial structure and physicochemical property distribution (see Fig. S5). It is also possible that, should this ionic/polar islet region be part of the epitope recognized by the antibody, the latter could pull N1158 away from the membrane to form an interface with itself, stabilizing the state we found to be transient for isolated FSL. D1153, another residue which was part of the hydrophilic islet, was poorly accessible in the pre-fusion state (Table 1), with only roughly 30% of the side chain exposed. Its accessibility was similar in the state tightly bound to the membrane, while in the loosely bound state it became much more accessible to the solvent as well as anti-1147 antibodies if they indeed recognize the pattern D1153 is part of.

*Phl p 7* interacts with the antibody via a number of hydrophilic residues in the disordered fragment downstream of the helix ending around D13: T14, N15 and G16. Obviously, there is not much point in comparing the conformations of these unstructured regions in peptide 1147 and *Phl p 7* because in the latter case antibody stabilizes one of multiple available states, which can exist for the free peptide. Such effects are well-known as so-called “induced-fit” interactions. At the same time, while we cannot draw a direct comparison between two disordered fragments, it is interesting that the counterpart of *Phl p 7’s* T14 in peptide 1147 is S1161 with a very similar polar side chain, and we do see states during MD in which it is positioned similarly in relation to the helix upstream, and we cannot exclude the possibility of this residue’s contribution to antibody binding. If this were the case, one would speculate that, were peptide 1147 to be extended to include P1162 and D1163 (kink-forming and polar, respectively), antibody binding might become more efficacious if a *Phl p 7-*like scenario does indeed occur. Also, of interest might be H1159, which faces the coiled-coil lumen in the pre-fusion state and can also hide itself away in the state tightly bound to the membrane, while appearing perfectly accessible in the loosely bound state. If this residue were part of the epitope recognized by human anti-1147 antibodies, this would also partially explain why the antibodies do not interact with the trimeric protein within the intact virus. Hydrophilic residue accessibility is even poorer in the post-fusion state. Overall, we could conceivably be looking at a state different from the two known stable ones of the full-sized trimeric protein in which the antibody recognizes the peptide 1147 region.

## 3. Conclusions

Predictions obtained using modeling tools imply that, under certain conditions, peptide 1147 would be capable of interacting with anti-1147. Experimental data, however, showed that, even at high concentrations, anti-1147 antibodies isolated from Covid patients lack the ability to block the virus. This fact would indicate that the conformational rearrangement of the stem region during cell infection most probably does not generate a conformation of the 1147 epitope recognized by, or accessible to anti-1147 antibodies. This means that therapeutic strategies based on the use of antibodies themselves or corresponding peptide vaccines will be ineffective.

## 4. Experimental

### Computer simulations

Topology files for the FSL construct and its control variant, in which acetylated cysteamine was attached to the spacer moiety in lieu of the peptide, were created using in-house software. On all occasions, they were aligned in such a manner as for the lipid tails to be parallel to the bilayer plane normal, and dihedrals were adjusted in order for the non-membrane-embedded part of the molecule, in its initial position, to also be oriented along the bilayer plane normal (similarly to the orientation shown in Fig.2).

On all occasions the molecule was inserted in the system so that the lipid tails would be fully submerged into the membrane. MD simulations were performed using the GROMACS 2020.4 package [17] and the CHARMM36 force field [18-22]. An integration time step of 2 fs was used and 3D periodic boundary conditions were imposed. The spherical cut-off function (12 Å) was applied to truncate van der Waals interactions. Electrostatic interactions were treated using the particle mesh Ewald (PME) method [23] (real space cutoff 12 and 1.2 Å grid with fourth-order spline interpolation). The TIP3P water model was used [24], and Na+ and Cl− ion parameters for counter ions were implemented. Simulations were performed at a temperature of 325 K and 1 bar pressure was maintained using the V-rescale [25, 26] algorithms with 0.5 and 5.0 ps relaxation parameters, respectively, and a compressibility of 4.5 × 10^−5^ to 10^-1^ bar for the barostat. The protein along with membrane lipids and solvent molecules were coupled separately. Semi-isotropic pressure coupling in the bilayer plane and along the membrane normal was used in the simulations. Before the production runs, all systems were minimized over 2000 steps using a conjugate gradients algorithm, followed by heating from 5K to 325 K over 50000 steps, during which internal coordinates of the protein and ligand heavy atoms were restrained. Production runs had lengths of 0.05 to 0.5 μs depending on the system. Bonds with an H atom were constrained via the LINCS algorithm [27].

All MD trajectories were processed using the *trjconv* utility from the GROMACS 2020.4 package to remove 3D periodic boundary conditions and to obtain an output frequency of 100 ps per frame. Coordinates were extracted using the *traj* utility, radii of gyration were calculated using the *gyrate* utility, while cluster analysis was performed via the *cluster* utility, all from the GROMACS package. Intermolecular contacts were calculated as described elsewhere [28]. Molecular hydrophobicity potential maps were created as described earlier [29]. Intermolecular interfaces were calculated using DSSP [30]. A residue was considered to be located on the interface if the difference between its ASA values in a single helix and in a helical oligomer was ≥ 25 Å^2^ (≥ 10 Å^2^ for glycine). Exposed and buried residues were identified using the NACCESS computer program [31]; a residue was considered exposed if the relative accessibility of its side chain was greater than or equal to 45%. Molecular editing and graphics rendering were performed using PyMOL v. 2.4.0 (The PyMOL Molecular Graphics System, Schrödinger, LLC) and UCSF Chimera [32].

### Sequence motif search

In order to identify complexes with antibodies potentially similar to those formed by peptide 1147, a search was run across the RCSB database (https://www.rcsb.org) for the motif “[ED][ED]x[ED][KR]xx” paired with the keyword “antibody” (where “x” corresponded to any amino acid except E/D/H/K/R/P). The complexes were then examined to select those in which the antibody was bound to the motif of interest as opposed to other part of the protein. Synthesis of FSLs is described elsewhere [5,7] using >95% purity peptides obtained from Synpel Chemical (Prague, Czech Republic).

### Synthesis of affinity adsorbent

2 ml of amino Sepharose 6FF (GE Healthcare, USA) was washed several times with 0.2 M aqueous NaHCO_3_, then with water, and, finally, with DMSO. A solution of 3-maleimidopropionic acid *N*-hydroxysuccinimide ester Mal-βAla-ONSu (23.4 mg, 88 micromoles) in DMSO (1 ml) was added to the amino Sepharose in DMSO, gently mixed and incubated at room temperature for 1.5 hrs. The gel was washed several times with DMSO+0.2% AcOH to remove excessive Mal-βAla-ONSu, then once with 2 ml of water. Peptide 1147 (4.35 mg, 1.71 µmol, as a TFA_5_ salt, according to TLC data, contains 5-7% of S-S dimer) was dissolved in water (0.5 ml) and immediately added to the gel suspension in water, gently mixed, 1 ml of 0.2 M NaOAc (the solution was adjusted to pH 6.4 by addition of AcOH) was added, and the gel was thoroughly mixed. The suspension quickly forms a foam during stirring, which disappears as the peptide reacts. After 20 min (TLC of the supernatant shows complete reaction of the peptide), mercaptoethanol solution was added to the suspension (88 micromoles, 344 µl of 20 mg/ml solution in water), mixed therewith and incubated for 20 min. The adsorbent was then washed several times with 0.2% AcOH, water, 0.2 M NaHCO_3_, water, and, finally, with 25% EtOH; it was stored in 25% EtOH at 4 °C. 1 ml of the resulting adsorbent 1147-Sepharose 6FF contains ∼ 0.8 micromoles of peptide 1147.

### Sera

Convalescent serum samples (National Medical Research Center for Obstetrics, Gynecology and Perinatology, Moscow, Russia) were obtained from female patients recovered from PCR -confirmed COVID-19 between May and June 2020. The study was approved by the local Ethics Committee (Approval number 4-23/04/2020); written informed consent was obtained from the subjects. Negative control serum samples were collected from healthy adult donors before 2019 showing no apparent signs of COVID at the time of sample collection and for which negative RT-PCR results were obtained (N.V. Sklifosovsky Research Institute of Emergency Care, Moscow, Russia). Group O erythrocytes were collected from healthy adult donors.

### Isolation of anti-1147 antibodies

Pooled blood serum from convalescent donors [5] was subjected to complement deactivation via incubation at 56 °C for 30 min and centrifuged at 7000g. The supernatant was diluted with phosphate-buffered saline (PBS, 0.15 M, pH 7.4) containing 0.02% NaN_3_ (PBS-Az) at a ratio of 1:3. A column with 1147-Sepharose 6FF (0.8 µmol of the peptide per 1 ml) as adsorbent was washed and equilibrated with PBS-Az at 25°C. After preparation, serum was passed through the column (at a serum-to-adsorbent ratio of 15:1) at 25 °C and at a flow rate 0.4 ml/min. The column was washed with five volumes of PBS-Az/0.1% Tween-20 followed by 20 volumes of PBS-Az. Bound antibodies were eluted with Tris-OH (0.2 M, pH 10.4) containing 0.02% NaN_3_ at a flow rate 0.2 ml/min. Fractions containing antibodies were immediately neutralized using glycine-HCl buffer (2M, pH 2.5) and pooled. The eluate from the 1147-Sepharose 6FF column was passed through a ligand-free Sepharose 6FF column equilibrated with Tris-OH/glycine-HCl (0.2 M, pH 7.4) containing 0.02% NaN_3_ at 25 °C at a flow rate of 0.2 ml/min to remove serum antibodies directed against affinity matrix. The material not retained on the column was collected, and concentration was evaluated by measuring absorbance at 280 nm (assuming 1% Ig absorbance at 280 nm to be 14.0) using a spectrophotometer (Ultrospec 3100 pro, Amersham, Buckinghamshire, UK). Antibodies were then concentrated in PBS (0.15 M, pH 7.4) without NaN_3_ using an Amicon Ultra Centrifugal Filter Unit, 10 kDa cutoff (Merck, Darmstadt, Germany) at 15 °C, 5500g. Antibodies were stored at 4 °C for several weeks or kept frozen at -20 °C when a longer storage period was required. Adsorbent ligand-free Sepharose 6FF was regenerated with NaOH (0.1 M) and verified as clean (UV control) before reuse.

### Artificial membrane-based EIA assays

#### One-stage assay

To each well of 96-well NUNC Maxisorp plates, 100 µl of an ethanol solution containing FSL-peptide (64, 32, 16, 8, 2 or 1 mol% from PC), PC (25 μg/ml), and cholesterol (50 μg/ml) was added. The wells were dried at 37 ^o^C. Plates were blocked with 3% BSA in PBS for 1 h at 37 ^o^C and washed three times with PBS containing 0.1% Tween-20 (wash buffer). Human plasma (in serial twofold dilutions starting from 1:20, 100 μl per well) was added, and the plates were incubated at 37 ^o^C for 1 h and washed three times with the wash buffer. Anti-human IgG-Biotin conjugate in PBS containing 0.3% BSA was added, and the plates were incubated at 37 ^o^C for 1 h and washed three times with the wash buffer. Plates were incubated with Str-HRPO conjugate (1:2000 in PBS containing 0.3% BSA) at 37 ^o^C for 1 h and washed three times with the wash buffer. Finally, the color was developed via 30 min incubation with 0.1 M sodium phosphate/0.1 M citric acid buffer, containing 0.04% *o*-phenylenediamine and 0.03% H_2_O_2_, and the reaction was stopped by adding 50 μL 1 M H_2_SO_4_. Absorbance was recorded at 492 nm in a Multiskan MCC/340 microtiter plate reader (Perkin Elmer, Turku, Finland). The control wells contained no FSL construct.

#### Two-stage assay

To each well of 96-well NUNC Maxisorp plates, 100 µl of an ethanol solution containing PC (25 μg/ml), and cholesterol (50 μg/ml) was added; the wells were dried at 37 ^o^C. Then peptide-FSL was added into the wells (100 μl per well), in PBS (pH 7.4) and incubated at 50 ^o^C for 1 h. The plates were blocked via incubation with 3% BSA in PBS at 37 ^o^C for 1 h and washed three times with PBS containing 0.1% Tween-20 (wash buffer). Human plasma (1:160; 100 μL per well) was added and the plates were incubated at 37 ^o^C for 1 h, washed three times with the wash buffer. Anti-human IgG-biotin conjugate in PBS was added; all the remaining steps were the same as for the one-stage method.

### Direct EIA

PolySorp microtiter plates (Nunc, ThermoFisher Scientific, Denmark) were coated with 5 µM (60 µl/well) peptide-FSL or PEG-FSL (as a background control) in PBS (0.15 M, pH 7.4) at 37 °C for 120 min and washed. The plates were blocked with 1% BSA in PBS (60 µl/well) at 37 °C for 45 min. Affinity-purified antibodies at a concentration of 1 µg/ml in PBS containing 0.3% BSA or blood sera diluted at a ratio of 1:5 in PBS containing 0.3% BSA were added (50 µl per well), incubated at 37 °C for 60 min and washed. Then HRP-labeled anti-human IgG (Invitrogen, USA, working dilution 1:12000) in PBS containing 0.3% BSA, 50 µl per well, was added to the plates and incubated at 37 °C for 60 min and washed. Color was developed via 20-min incubation at room temperature in 0.1 M sodium phosphate/0.1 M citrate buffer containing 0.04% of *o*-phenylenediamine and 0.03% of H_2_O_2_. The color reaction was stopped by adding 1M H_2_SO_4_. The absorbance was read at 492 nm with a multitask plate reader (Wallac1420 Multilabel Counter, Victor^2^, Perkin Elmer Life Sciences, Finland). Between stages the plates were washed four times with PBS containing 0.1% Tween-20.

All the tests were performed at least in duplicate; the differences between readings (intra-assay) did not exceed 5%. The data obtained were processed using Microsoft Excel.

### Virus neutralization potency of anti-1147 antibodies in cell culture system

To study the virus neutralizing activity (VNA) of the blood sera of convalescents and affinity-isolated antibodies, the Vero C1008 cell culture line (cell monolayer in a flask) and SARS-CoV-2 coronavirus (variant B) were used. Cells were cultivated using Medium 199 with Hanks’ salts, supplemented with 7.5% fetal bovine serum (FBS), glutamine, glucose, and gentamicin. The working dilution of the virus was prepared in Hanks’ solution with 2% FBS and 100 U/ml of streptomycin sulfate and benzylpenicillin by making serial 10-fold dilutions of the initial virus-containing suspension until the final concentration of the virus in the sample was 200 BAU per ml. The VNA was determined using a neutralization assay; the metric used was the suppression of negative colony formation caused by SARS-CoV-2 virus in a one-day monolayer of Vero C1008 cells under agar overlay. A mixture of equal volumes of the sample tested and SARS-CoV-2 virus culture at a working dilution was incubated at a temperature between 36.5 and 37.5 ^o^C for 60 min; 0.5 ml of the mixture was then applied onto a monolayer of Vero C1008 cells. After 60 min, the inoculum was decanted; the primary agar overlay for SARS-CoV-2 virus was then applied, and the monolayer was incubated at 36.5 to 37.5 ^o^C for two more days. Next, a secondary staining agar overlay with 0.1% Neutral Red was applied onto the infected cell monolayer and incubated at 36.5 to 37.5 ^°^C for 24 hours; at this stage, negative colonies were counted. The highest dilution of serum that caused a 50% reduction in the number of plaques, compared to the average viral control, was considered as the VNA titer.

## Abbreviations

Ab: antibody
MD: molecular dynamics
FSL: Function-Spacer-Lipid construct
Ig: immunoglobulin
MHP: molecular hydrophobicity potential
POPC: 1-palmitoyl-2-oleoyl-glycero-3-phosphocholine
Chol: cholesterol
DOPE: 1,2-dioleoyl-sn-glycero-3-phosphoethanolamine
peptide 1147: amino acid residues 1147-1161 (SFKEELDKYFKNHTS) of the SARS-Cov-2 protein S

## Author contributions: CRediT

Elena Aliper – Investigation, Writing – original draft, Writing – review and editing

Ivan Ryzhov – Investigation, Writing – review and editing

Polina Obukhova – Investigation, Writing – review and editing

Alexander Tuzikov – Investigation, Writing – review and editing

Oxana Galanina – Investigation, Writing – original draft

Marina Ziganshina – Writing – original draft, Writing – review and editing

Gennady Sukhikh – Writing – review and editing

Nikolay Krylov – Investigation, Writing – review and editing

Stephen Henry – Writing – review and editing

Roman Efremov – Project administration, Supervision, Writing – original draft, Writing – review and editing

Nicolai Bovin – Project administration, Supervision, Writing – original draft, Writing – review and editing

## Data availability

No data was used for the research described in the article.

