## Supplementary Data for "Immune Antibodies Recognizing the Stem Region of SARS-CoV-2 Spike Protein: Molecular Modelling and *In Vitro* Study of Synthetic Peptides Presentation to the Antibodies"

### Table of contents


| | |
|---|---|
| Molecular dynamics simulation | 3 |
| Figure S1 | 5 |
| Figure S2 | 6 |
| Figure S3 | 7 |
| Figure S4 | 8 |
| Figure S5 | 9 |
| Figure S6 | 11 |

Molecular dynamics simulation

One monolipidated FSL molecule (Fig. 1) was inserted in a bilayer made up of 1-palmitoyl-2-oleoyl-glycero-3-phosphocholine (POPC) and cholesterol (Chol) at a ratio of 2:1 in order to mimic the lipid membrane. As a control, the cysteine moiety attached to the peptide was replaced with acetylated cysteamine, in order to verify whether the construct without the peptide was in any way membrane-active. Initially, 10 runs each 0.05 μs long were performed for the FSL in the bilayer, as is commonly done to capture the membrane binding mode of membrane-active proteins. Of these three (runs 1, 3 and 4) were extended to a total of 0.5 μs. Two 0.5-μs-long runs were performed for the "FSL minus peptide" control.

As can be seen from the dynamics of the Z-coordinate of the S atom in the linking Cys of the FSL and in the Ac-cysteamine of the control (Fig. S2), the former demonstrated a certain capacity for staying close to the water-lipid interface, the distance between the S atom and the center of mass of the P atoms of the upper leaflet amounting to 1.14 ± 0.89 nm, 0.85 ± 0.57 nm and 1.91 ± 0.92 nm in runs 1, 3 and 4, respectively. Meanwhile the control molecule behaved in a more random way, seemingly having no preference for the interface as opposed to the aqueous environment, and demonstrated the same parameter of 1.60 ± 1.31 nm and 1.80 ± 1.16nm in the two replicas. This was further corroborated by the analysis of radius of gyration (Fig. S3) across the trajectories where its value drops to some extent in the case of the FSL, with mean values being 0.87 ± 0.16 nm, 0.94 ± 0.18nm and 1.01 ± 0.21 nm in run 1, run 3 and run 4, respectively. The effect is more pronounced in runs where the peptide binds to the membrane (200 ns to 300 ns in run 1, from approx. 100 ns onwards in run 3), whilst in the linker control it tended to revert to values close to the maximum observed and averaged 1.08 ± 0.21 nm and 1.08 ± 0.22 nm in the two runs.

The dynamic of the peptide's secondary structure in the three extended runs varied. In run 1, the initial helical conformation disintegrated, while in run 3, in which the peptide ended up buried in the membrane, and in run 4, in which it failed to do so, the initial α-helix was well preserved, especially up to residues 1555-1556 (Fig. S4).

In run 1, only one relatively long-lived cluster was identified corresponding to a membrane bound state, from 220.5 to 283.5 ns (120 out of 127 frames over the specified time interval), with the RMSD cutoff value of 0.15 nm. In this state the disordered peptide interacts with the membrane via its N-terminus, in agreement with data on non-covalent protein-lipid interactions. Indeed, residues contributing over the greatest number of frames appear to be F1148 and K1149 with 61% and 54% of the total number of frames (Fig. S4).

In run 3, the cutoff value of 0.15 nm only yielded one major cluster, while the cutoff value of 0.1 nm allowed some further differentiation. The most notable clusters turned out to exist at 39.5 to 354 ns (96% of frames over the interval) and at 354 to 500 ns (nearly 100% frames over the interval). The majority of the residues interacted with the membrane lipids for almost the entire duration of the trajectory, except the C-terminus, which engaged in non-covalent interactions to a lesser extent. We were also able to identify a *bona fide* binding mode for peptide 1147 within the FSL (Fig. S5 A-B). It appears that peptide 1147 is located on the water-lipid interface, lying more or less parallel. The hydrophobic surface of the peptide faces away from the interface, while the hydrophilic residues are exposed to the aqueous environment. The submerged hydrophobic patch is made up by F1148, L1152, Y1155 and F1156. Additionally, the charged K1149 also faces down, its amino group forming long-lived hydrogen bonds and salt bridges with nearby phosphate groups of POPC, making it conceivably inaccessible to antibody. In run 3, shortly before cluster 1 forms and peptide 1147 delves deeper into the membrane, it associates with what could be described a "loosely bound state" characterized by only major hydrophobic moieties (and K1149) and becomes spatially inaccessible to any potential ligands. Peptide/lipid interfaces in model FSL constructs can thus be compared against protein/protein ones in different states of the protein S trimer (summarized in Fig. 7).

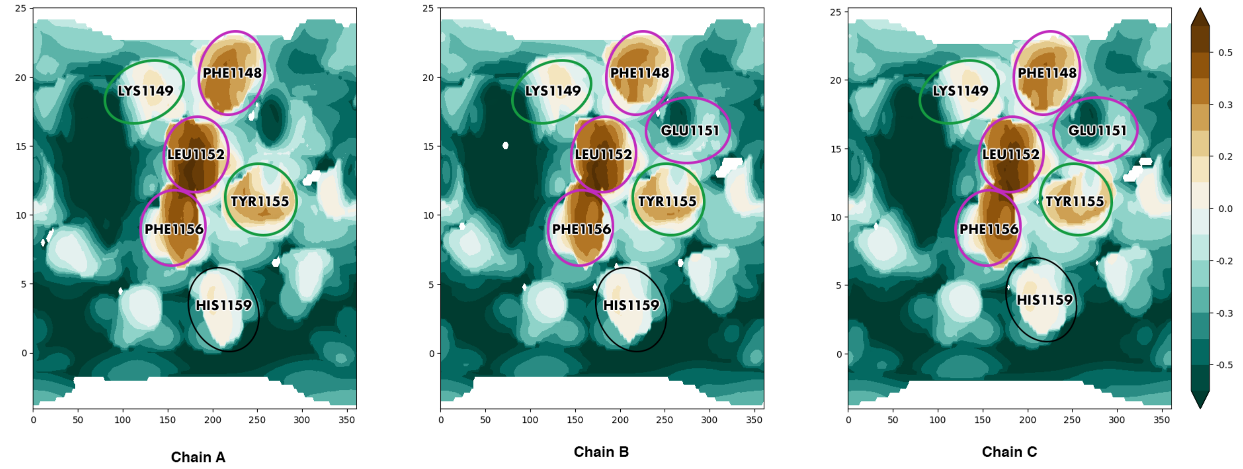


**Figure S1. Helix-helix interfaces for peptide1147 in the pre-fusion state of SARS-CoV-2 spike protein (PDB ID 6XR8 [10]).** Cylindrical projection of surface MHP (molecular hydrophobicity potential) distribution is used. Axis values correspond to the rotation angle around the helical axis and the coordinate along the latter (Å), respectively. An MHP scale (in logP octanol-1/water units) is presented on the right. The maps are colored in accordance with the MHP values [11], from teal (hydrophilic areas) to brown (hydrophobic ones). Identical positions across genus *Betacoronavirus* are encircled in purple, conservative and semi-conservative residues are encircled in green, non-conservative residues present on the helix/helix interface are encircled in black.

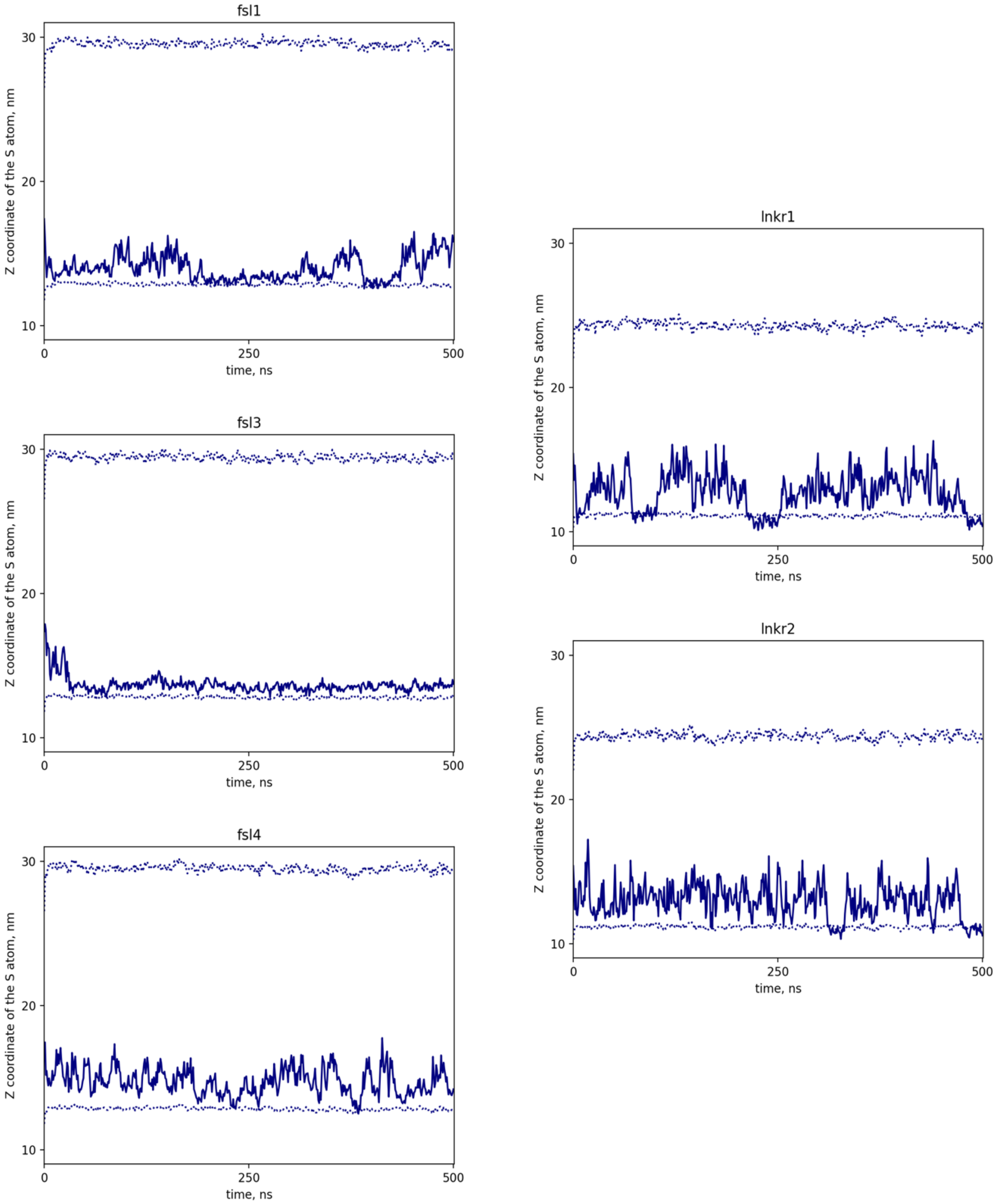


**Figure S2. Z-coordinate of the sulfur atom in the Cysteine of the Function-Spacer-Lipid (FSL) and in the Ac-cysteamine of the control molecule (lnkr) in MD simulations.** The Z-coordinate of the sulfur atom is shown as a solid line, while those of the centers of mass of phosphate groups of the upper leaflet (below) and the periodic image of the lower leaflet (above) of the membrane are shown as dotted lines. The zone between two dotted lines corresponds to the aqueous environment.

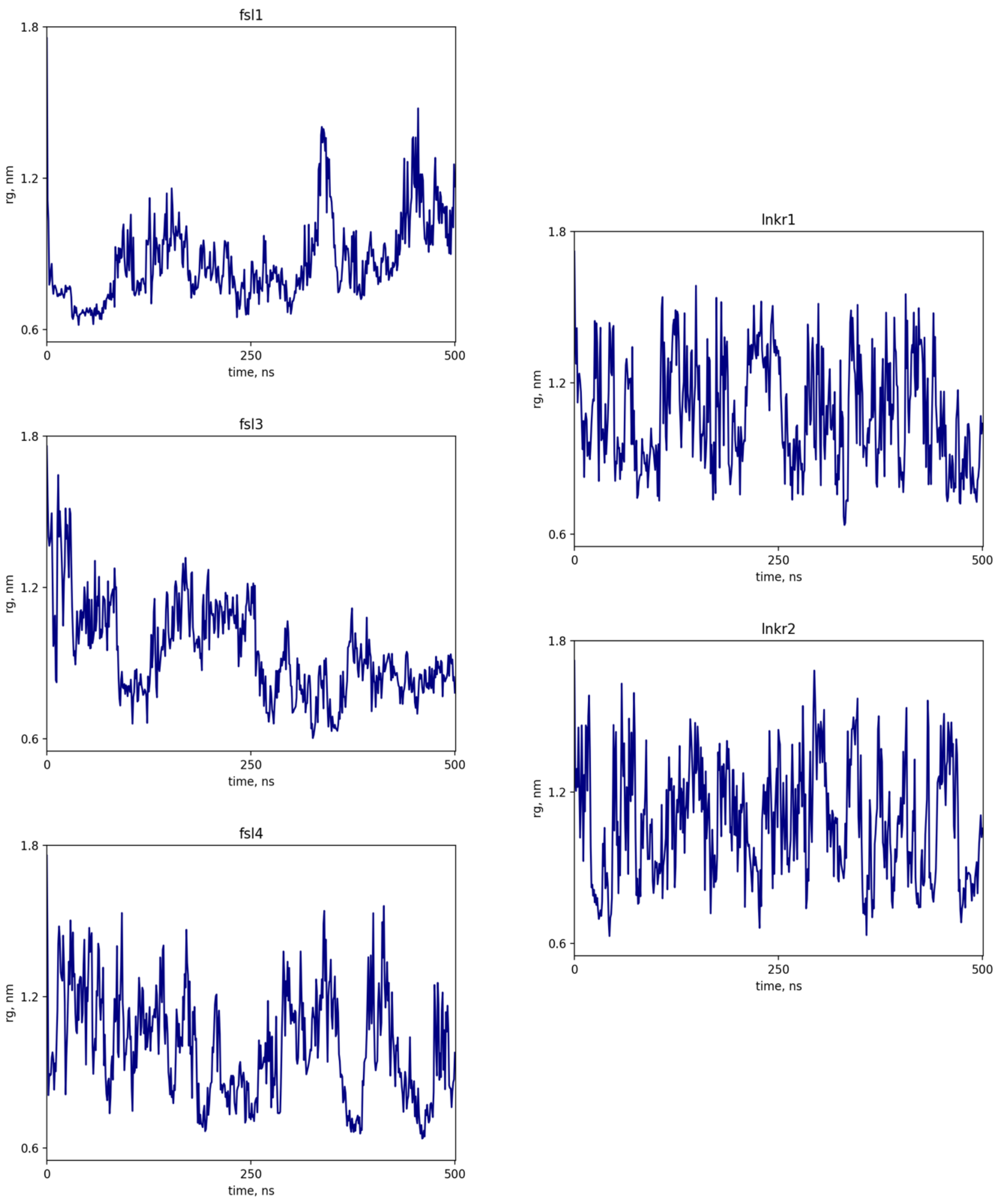


**Figure S3. Radius of gyration of the linker of the FSL and of the control molecule (lnkr) in MD simulations.**

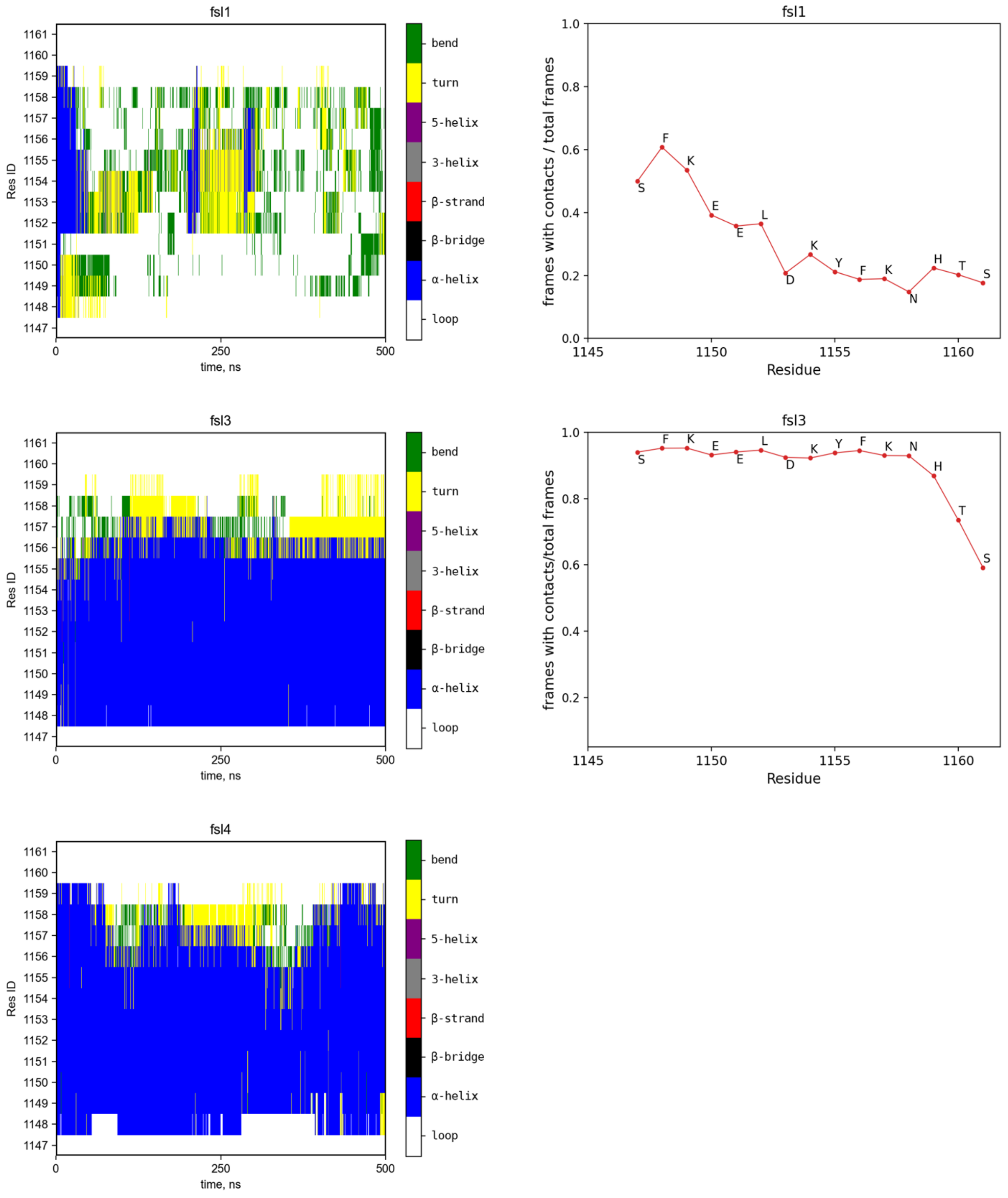


**Figure S4. Secondary structure of the peptide within the FSL in run 1 (top left), run 3 (middle left) and run 4 (bottom left); contribution to non-covalent interactions on the part of peptide 1147 residues in run 1 (top right) and run 3 (middle right).**

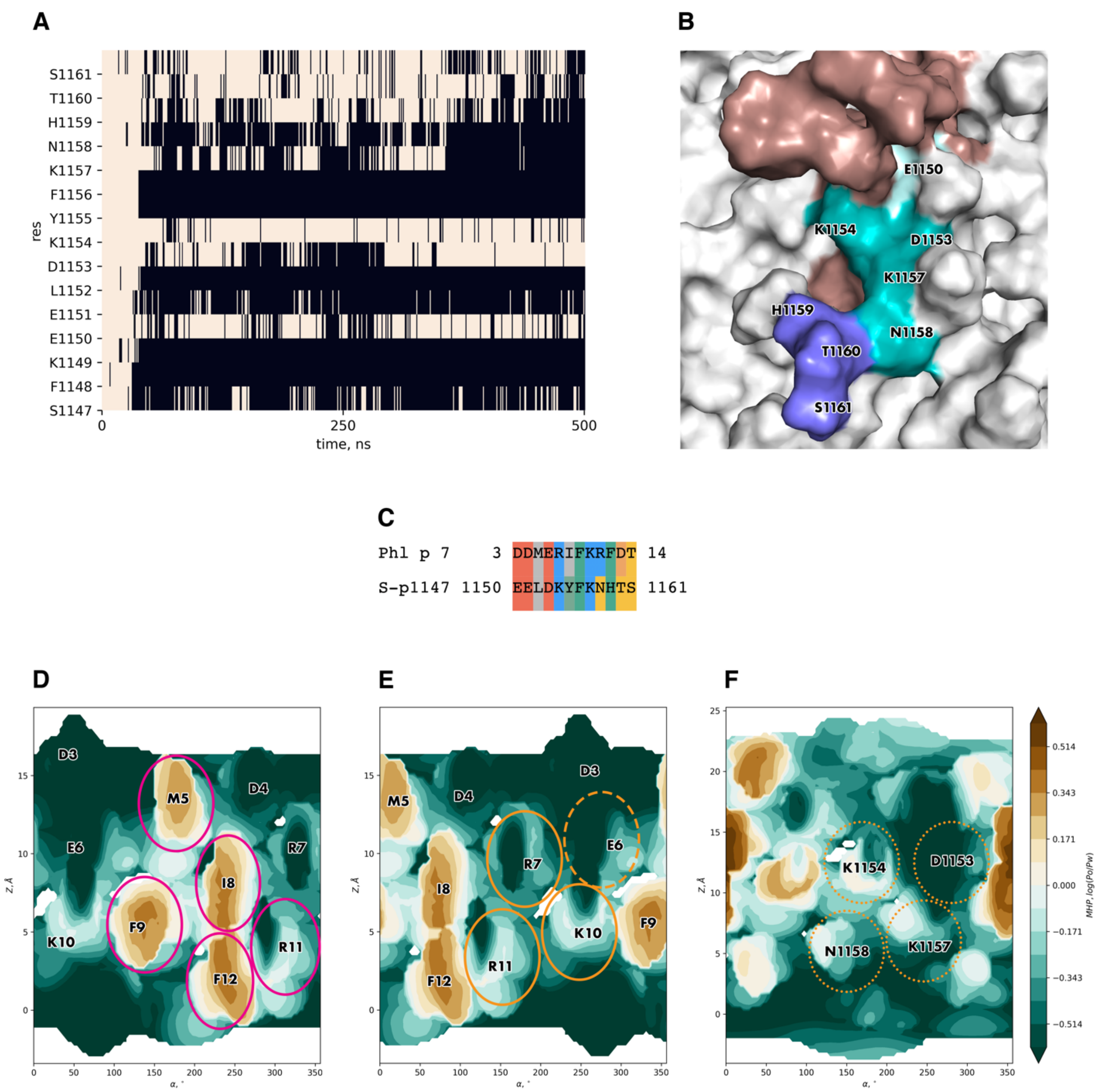


**Figure S5. The binding mode identified for peptide 1147.** (A) Heatmap representing accessible surface area for each residue across the trajectory. The membrane-buried state and exposed state are shown as black and off-white, respectively. (B) A 3D view of the membrane surface with anchored FSL from above. The FSL is colored brown, while the pattern of polar residues similar to that in Phl p 7 is colored teal, with E1150, another solvent-accessible polar residue, colored pale cyan, and the C-terminal hydrophilic fragment (1159-1161) colored purple blue. (C) Sequence comparison between peptide 1147 and *Phl p 7*. Negatively charged, positively charged, polar, hydrophobic and aromatic residues are highlighted in red, blue, yellow, grey and green, respectively. Y1155 in peptide 1147 is highlighted in greyish green to indicate it is both aromatic and hydrophobic and D13 in *Phl p 7* is highlighted in golden orange to indicate it is both negatively charged and polar; in both cases this is done to accentuate the similarity between the two sequences in these positions. (D) MHP distribution on the surface of *Phl p 7* helix 1 contacting other parts of the allergen molecule. Residues in the peptide/peptide interface are encircled in pink. (E) MHP distribution on the surface of *Phl p 7* helix 1 contacting its antibody. Residues in the peptide/antibody interface are encircled in golden, while a residue not corresponding to the interface criteria but interacting non-covalently with the antibody is enclosed in a golden dashed line. (F) MHP distribution on the surface of peptide 1147 exposed to the aqueous environment modelled as part of the FSL. Residues found in positions corresponding to those in Phl p 7 in accordance with the alignment in section C are encircled in golden dotted lines. See legend to Fig. S1 for more detail on MHP mapping.

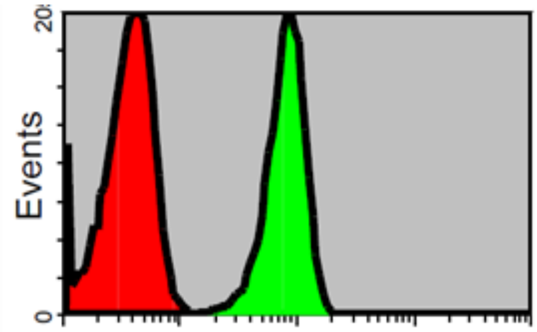


**Figure S6. Example of binding of Covid-19 positive (in green) antibodies to FSL-1147 when inserted into blood group O erythrocytes. The red peak is the negative control (PBS).**
